# Fission yeast Pdk1 kinase regulates cytokinesis and eisosomes

**DOI:** 10.1101/2025.07.16.665175

**Authors:** Madeline L. Chrupcala, Mackenzie J. Flynn, James B. Moseley

## Abstract

The conserved phosphoinositide-dependent protein kinase PDK1 regulates cell growth and stress signaling in eukaryotes. In the fission yeast *S. pombe*, Pdk1 has been linked to cytokinesis, which could point to new functions for this kinase family. Here, we discovered that Pdk1 localizes to eisosomes, which create invaginations in the plasma membrane, in addition to the spindle pole body (SPB). Pdk1 promotes phosphorylation of the core eisosome protein Pil1 and regulates eisosome length. Dysregulated eisosomes are not responsible for cytokinesis defects previously observed in *pdk1Δ* cells. Instead, we found that Pdk1 regulates localization of the anillin-like protein Mid1 and the protein kinase Sid2, which promotes cytokinesis as part of the septation initiation network (SIN). Our combined results provide insights into the role of Pdk1 in eisosomes and cytokinesis, which extend the functions of this conserved protein kinase family beyond canonical growth control pathways.

## INTRODUCTION

Conserved signal transduction pathways coordinate cell growth and division in eukaryotic cells. The phosphoinositide-dependent protein kinase PDK1 functions as a master regulator of cell growth pathways in organisms from yeast through humans (Pearce *et al*., 2010; Leroux *et al*., 2018). PDK1 is primarily known for its role in phosphorylating the activation loop of more than 23 downstream AGC family protein kinases in response to growth factor signals. These targets include kinases involved in cell proliferation, migration, and metabolism such as Akt, PKC, ribosomal S6 kinase, and p70 S6 kinase (Mora *et al*., 2004; Bayascas, 2010; Leroux and Biondi, 2023). Underscoring its critical role in conserved growth pathways, PDK1 knockout mice die at embryonic day E9.5 with severe developmental defects (Lawlor *et al*., 2002). In addition, PDK1 overexpression has been observed in many human cancers and is associated with poor clinical outcomes (Pearn *et al*., 2007; Eser *et al*., 2013; Zabkiewicz *et al*., 2014; Du *et al*., 2016).

Studying PDK1-related kinases in diverse organisms has the potential to reveal new functions beyond activation of AGC kinases. The fission yeast *Schizosaccharomyces pombe* expresses two PDK1-related kinases called Ksg1 and Pdk1. Similar to human PDK1, Ksg1 has an amino terminal kinase domain and a carboxyl terminal pleckstrin homology (PH) domain, which can interact with lipids (Niederberger and Schweingruber, 1999) (Supplementary Figure S1A). *ksg1+* is an essential gene, and the Ksg1 protein appears to activate AGC kinases involved in functions including meiosis and cell wall biogenesis (Niederberger and Schweingruber, 1999; Gräub *et al*., 2003; Cipak *et al*., 2024). In contrast, far less is known about the non-essential *pdk1+* gene. *pdk1Δ* phenotypes include defects in cytokinesis and mitosis (Bimbó *et al*., 2005; Liu *et al*., 2021), which might suggest new functions for PDK1-related signaling. The Pdk1 protein sequence includes an amino terminal kinase domain but lacks a PH domain, similar to its budding yeast orthologs Pkh1 and Pkh2 (Casamayor *et al*., 1999). Budding yeast Phk1 and Phk2 localize to eisosomes (Roelants *et al*., 2002; Walther *et al*., 2007), which are invaginations at the plasma membrane. This localization is strongest for Pkh2, which regulates phosphorylation of the eisosome protein Pil1 (Roelants *et al*., 2002; Walther *et al*., 2007; Fröhlich *et al*., 2009; Paine *et al*., 2023). Pkh1 and Pkh2 have been linked to numerous functions including lipid signaling, cell wall regulation, and endocytosis (Friant *et al*., 2001; deHart *et al*., 2002; Roelants *et al*., 2002).

In this study, we investigated fission yeast Pdk1 with the goal of learning about new functions. We found that Pdk1 localizes to fission yeast eisosomes and promotes phosphorylation of the core eisosome protein Pil1 for proper eisosome formation. Septation defects in *pdk1Δ* cells are not due to dysregulated eisosomes, but rather to delayed assembly of the cytokinetic actomyosin ring. Our findings provide insight into the functions of a conserved protein kinase in a powerful model organism.

## RESULTS AND DISCUSSION

### Pdk1 regulates eisosomes

As a first step to study Pdk1 function, we examined the localization of Pdk1-mNeonGreen (mNG) in live cells. A previous study on fixed cells reported mitosis-specific localization of Pdk1 to the spindle pole body (SPB) and to a cortical band reminiscent of the assembling cytokinetic actomyosin ring (Bimbó *et al*., 2005). We observed prominent localization of Pdk1-mNG to the cell cortex in all cells, along with localization to the SPB (Figure 1A). Cortical Pdk1-mNG did not colocalize with Rlc1-mCherry, a marker of the assembling and mature cytokinetic ring (Figures 1B and S1B). We conclude that Pdk1 is not a component of the cytokinetic ring precursors, and instead localizes to other structures at the cell cortex.

**Figure 1.**
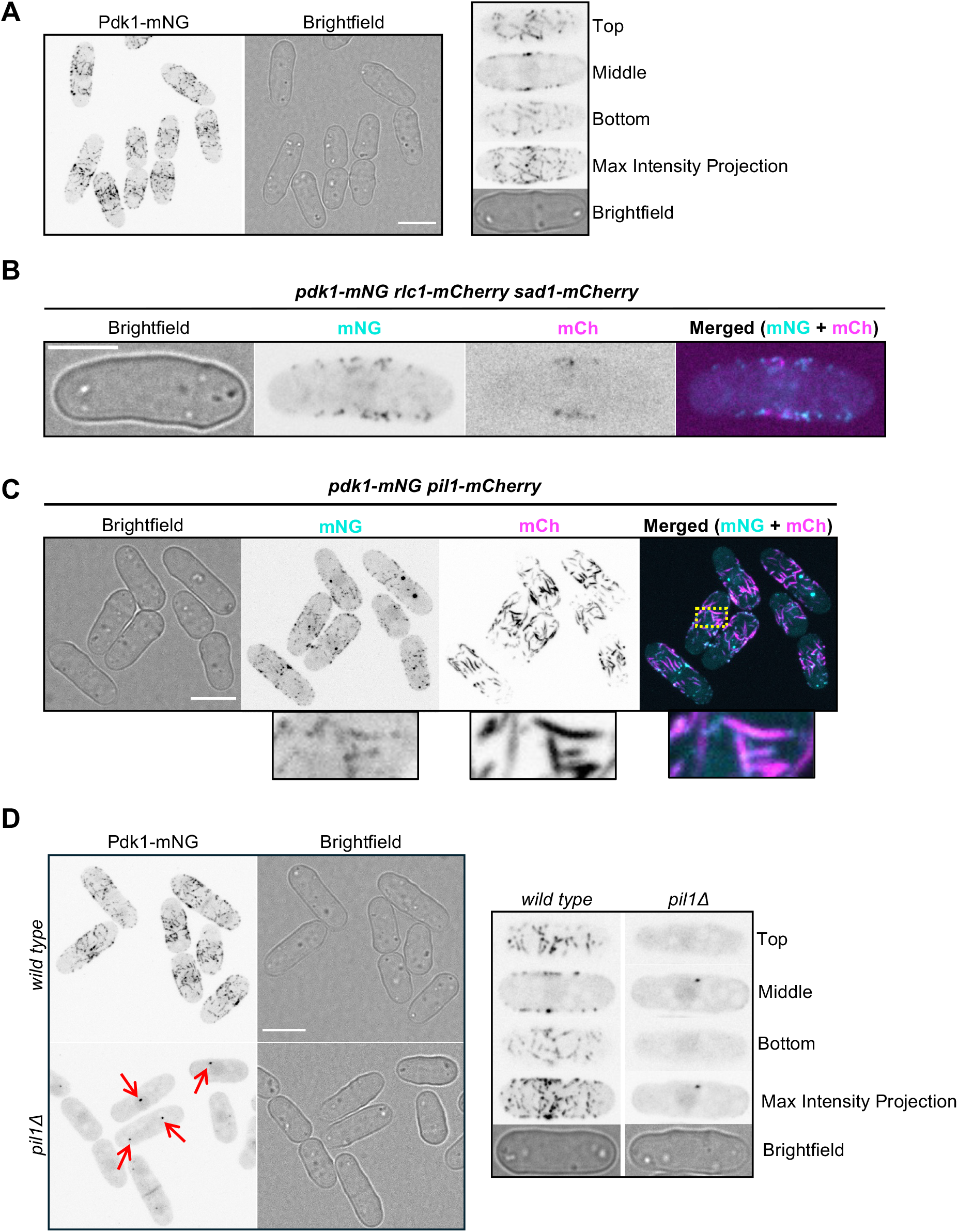
Pdk1 localizes to eisosomes. (A) Representative images of cells expressing Pdk1-mNG. Left, mNG images are maximum intensity projections with inverted fluorescence. Scale bar, 7 μm. Right, single focal planes and maximum intensity projection for a single cell. (B) Representative images of cells expressing Pdk1-mNG, Rlc1-mCherry to mark the cytokinetic ring, and Sad1-mCherry to label the Spindle Pole Body (SPB). mNG and mCh images are maximum intensity projections through the top half of the cell. Merged images are maximum intensity projections of mNG (cyan) and mCh (magenta) channels overlaid. Scale bar, 5 μm. (C) Representative images of cells expressing Pdk1-mNG and Pil1-mCherry to mark eisosomes. mCh and mNG images are maximum intensity projections with inverted fluorescence. Merged images are maximum intensity projections of mCh (magenta) and mNG (cyan) channels overlaid. Yellow boxed region is expanded at bosom. Scale bar, 7 μm. (D) Left, representative images of wild type and *pil1Δ* cells expressing Pdk1-mNG. mNG images are maximum intensity projections with inverted fluorescence. Scale bar, 7 μm. Red arrows mark SPBs. Right, single focal planes and maximum intensity projection for a single wild type or *pil1Δ* cell expressing Pdk1-mNG.

Cortical Pdk1 forms faint linear structures reminiscent of eisosomes, which are linear invaginations of the plasma membrane formed by the BAR domain protein Pil1 (Kabeche *et al*., 2011). This possibility is supported by the localization of budding yeast PDK1-related kinases to eisosomes (Walther *et al*., 2007; Fröhlich *et al*., 2009; Paine *et al*., 2023). We found that Pdk1-mNG colocalized with Pil1-mCherry (Figure 1C), even though the level of Pdk1-mNG cortical filaments was reduced by the Pil1-mCherry tag. We also found that Pdk1-mNG was absent from the cell cortex in *pil1Δ* cells, which lack eisosomes, and instead Pdk1-mNG was localized to the SPB and diffusively in the nucleus and cytoplasm (Figure 1D). We note that the role of Pil1 in recruiting Pdk1 to eisosomes may be indirect because all known eisosome proteins and associated membrane invaginations are lost from these structures in *pil1Δ* cells (Kabeche *et al*., 2011, 2015). These data show that the primary sites of Pdk1 localization in fission yeast cells are the SPB and eisosomes.

We next tested if Pdk1 regulates eisosomes. In budding yeast, conflicting reports have suggested that Pkh1/2 either promote or inhibit eisosome formation (Walther *et al*., 2007; Luo *et al*., 2008). To test this, we compared eisosome length in wild type and *pdk1Δ* cells. We found that Pil1-mCherry filaments were longer in *pdk1Δ* cells compared to wild type (Figure 2A-B). Next, we examined the effect of increased Pdk1 levels on eisosome length by generating a strain that expresses two copies of *pdk1+ (2x-pdk1)*. Pil1-mCherry filaments were shorter in *2x-pdk1* cells compared to wild type (Figure 2A-B). This supports a model where Pdk1 limits eisosome assembly by Pil1.

**Figure 2.**
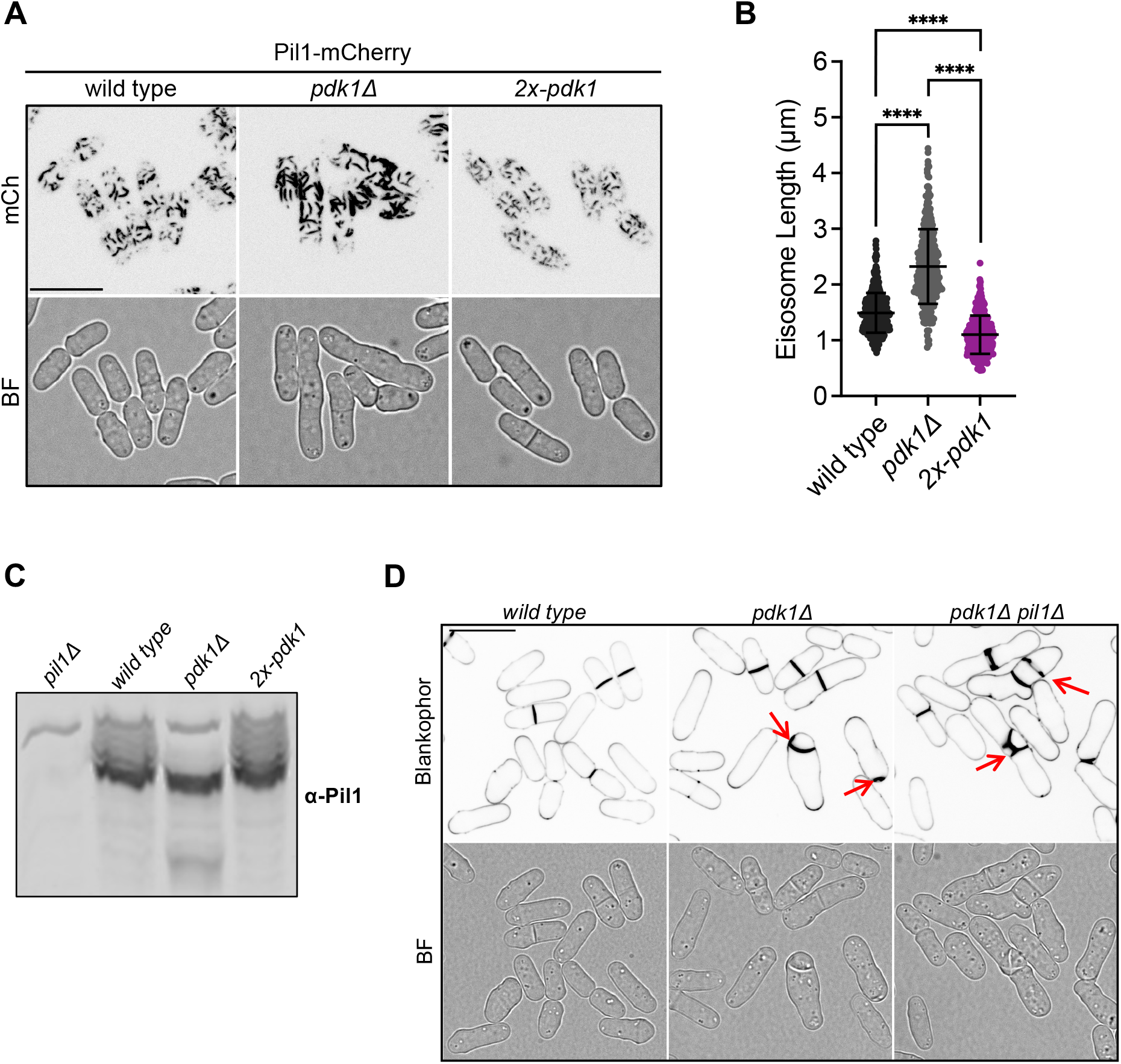
Pdk1 regulates eisosomes and promotes phosphorylation of Pil1. (A) Representative images of wild type, *pdk1Δ*, and *2x-pdk1* cells expressing Pil1-mCherry. mCh images are maximum intensity projections with inverted fluorescence. Scale bar, 14 μm. (B) Quantification of Pil1-mCherry eisosome length in the indicated strains. >235 cells were analyzed for each strain. Lines denote mean and standard deviation. Welch’s unpaired t-test was used to compare the two strains. **** indicates a *p* value < 0.0001. (C) Western blot analysis of Pil1 phosphorylation status in whole-cell lysates from *pil1Δ*, wild type, *pdk1Δ*, and *2x-pdk1* cells. (D) Representative images of cells stained with Blankophor to mark the cell wall for the indicated strains. Red arrows point to aberrant septa. Blankophor images are a single middle focal plane with inverted fluorescence. Scale bar, 14 μm.

To investigate the underlying mechanism, we tested the possibility that Pdk1 regulates Pil1 phosphorylation. Using Phos-bind SDS-PAGE, which slows the migration of phosphorylated proteins, we observed pronounced smearing of bands for Pil1 in wild type cells (Figure 2C). In contrast, Pil1 collapsed to a single band in *pdk1Δ* cells, showing loss of phosphorylation, and Pil1 phosphorylation was enhanced in *2x-pdk1* cells. We conclude that Pdk1 promotes Pil1 phosphorylation and limits eisosome assembly. Our combined results show that Pdk1 and Pil1 reciprocally regulate each other: Pil1 promotes Pdk1 localization to eisosomes, and Pdk1 limits the elongation of eisosomes.

Finally, we tested if dysregulated Pil1 causes the previously described septation defect in *pdk1Δ* cells. Eisosomes are known to be cleared from the cell division site prior to septation (Kabeche *et al*., 2011). If the septation defects of *pdk1Δ* cells are caused by excessive eisosomes, then eisosome removal by *pil1Δ* should suppress this defect. We tested this possibility by generating *pdk1Δ pil1Δ* double mutants. However, we found that abnormal septa remained present in *pdk1Δ pil1Δ* cells (Figure 2D), indicating that septation defects in *pdk1Δ* cells are not due to excess eisosomes. Therefore, we decided to examine cytokinesis in *pdk1Δ* cells.

### Cytokinesis defects in pdk1Δ cells

Fission yeast cytokinesis occurs in three defined and reproducible steps. First, cytokinesis proteins assemble at a series of cortical precursor structures called nodes, which are anchored by the anillin-like protein Mid1 and the protein kinase Cdr2 (Paoletti and Chang, 2000; Wu *et al*., 2006; Almonacid *et al*., 2009). Nodes condense to form the cytokinetic ring. Second, additional cytokinesis proteins are recruited to the ring in a step termed “maturation” (Wu *et al*., 2003, 2006; Pollard and Wu, 2010). Third, the cytokinetic ring constricts to separate the two dividing cells. To examine the role of Pdk1 in cytokinesis, we monitored wild type and *pdk1Δ* cells by timelapse microscopy using Rlc1-mNG to mark the cytokinetic ring and Sad1-mEGFP to mark the SPBs, which provide a temporal clock for assembly and constriction of cytokinetic events (Wu *et al*., 2003). Compared to wild type cells, the assembly of the cytokinetic ring was significantly slower in *pdk1Δ* cells (Figure 3A-B). The duration of ring maturation and constriction were unaffected in *pdk1Δ* cells (Figure 3C-D). Notably, cytokinetic rings failed to fully assemble and then fell apart without constricting in approximately 10% of *pdk1Δ* cells (Figure 3A). Our results identify ring assembly as a stage of cytokinesis that is defective in *pdk1Δ* cells.

**Figure 3.**
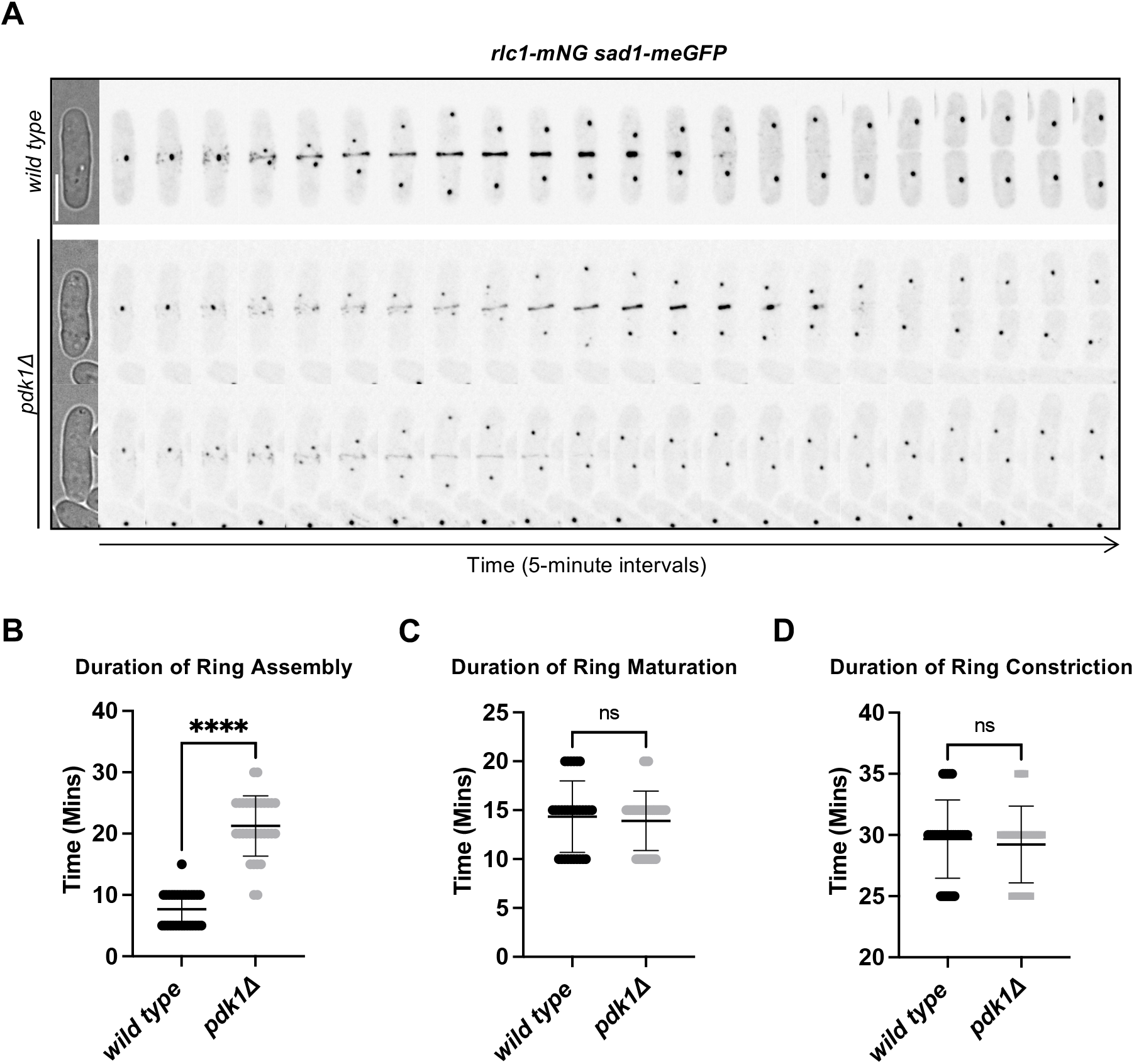
Loss of Pdk1 slows assembly of the cytokinetic actomyosin ring. (A) Representative time-lapse montage of wild type and *pdk1*Δ cells expressing Rlc1-mNG to mark the CAR and Sad1-meGFP to label the SPB. Images were captured in 5-minute intervals. Scale bar, 7 μm. (B-D) Duration of ring assembly (B), ring maturation (C), and ring constriction (D) in wild type and *pdk1*! cells. ti ≥ 30 cells each. Lines denote mean and standard deviation. All statistical tests in this figure are Welch’s unpaired t-tests. **** indicates *p* value < 0.0001, ns indicates *p* value > 0.05. The Materials and Methods section includes a full description of how each stage was defined and measured.

Our results suggest that *pdk1Δ* cells have defects in cytokinetic ring assembly. To extend this idea, we examined genetic interactions between *pdk1Δ* and the cytokinetic ring proteins Cdc15 and Rng2. Both Cdc15 and Rng2 make multiple protein-protein interactions within the cytokinetic ring, and Cdc15 also binds directly to lipids (Roberts-Galbraith *et al*., 2010; Laporte *et al*., 2011; Padmanabhan *et al*., 2011; McDonald *et al*., 2015; Ren *et al*., 2015). These proteins are essential for cytokinetic ring formation and cell viability, and the *cdc15-140* and *rng2-D15* mutations are temperature-sensitive (Nurse *et al*., 1976; Fankhauser *et al*., 1995; Eng *et al*., 1998). We observed negative genetic interactions between *pdk1Δ* and either *cdc15-140* or *rng2-D15* mutations (Supplementary Figure S1C-E). We found reduced growth and increased septation defects for *pdk1Δ cdc15-140* and *pdk1Δ rng2-D15* double mutants, when compared to the single mutants (Supplementary Figure S1C-E). These results reinforce the functional importance of Pdk1 in formation of the cytokinetic ring.

To gain mechanistic insight into cytokinesis defects in *pdk1Δ* cells, we examined the localization of key cytokinesis proteins in the absence of Pdk1. The cytokinetic ring assembles from precursor nodes, which are positioned by Cdr2 and Mid1. We did not observe defects in the distribution of Cdr2 nodes in *pdk1Δ* cells (Figure 4A). This result was confirmed by measuring the spread of Cdr2 nodes along the length of the cell, which was similar in wild type and *pdk1Δ* cells (Figure 4B). However, we found that Mid1 cortical nodes were more spread out in *pdk1Δ* cells compared to wild type cells (Figure 4C-D). This suggests that the disorganization of cortical Mid1 nodes may contribute to delayed cytokinetic ring assembly in *pdk1Δ* cells. We also tested if cytokinetic rings in *pdk1Δ* have altered levels of key proteins. We did not observe reductions in the levels of myosin proteins Rlc1, Myp2, and Myo2, formin Cdc12, or Rho-GAP Rga7 at the cytokinetic rings of *pdk1Δ* cells, although we note statistically significant increases for Myp2 and Cdc12 (Supplementary Figure S2A-E).

**Figure 4.**
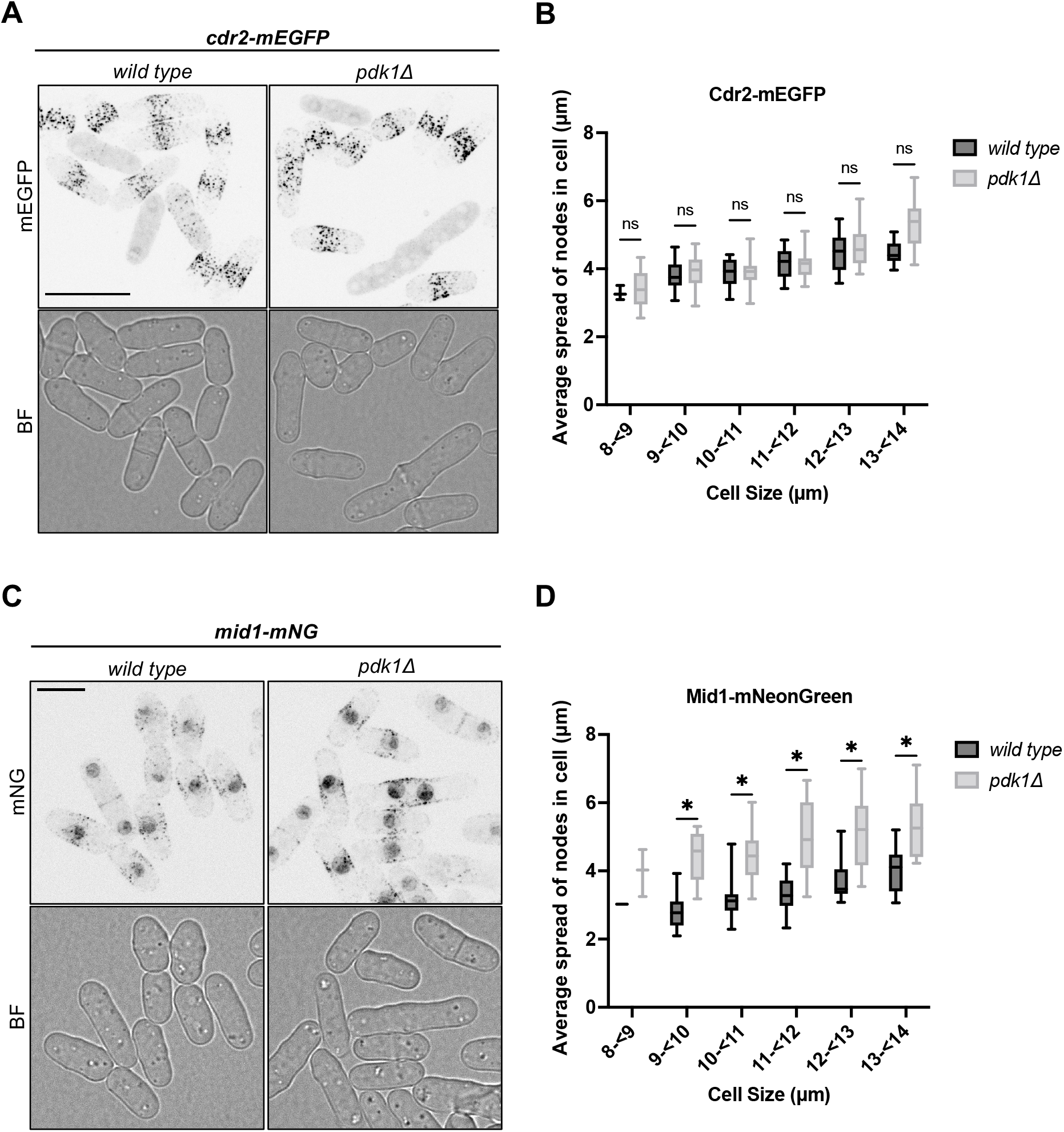
Disorganization of Mid1 cortical localization in *pdk1Δ* cells. (A) Representative images of wild type and *pdk1Δ* cells expressing Cdr2-mEGFP. mEGFP images are maximum intensity projections through the top half of the cell with inverted fluorescence. Scale bar, 14 μm. (B) Quantification of average Cdr2-mEGFP node distance versus cell size in wild type and *pdk1*Δ cells. Error bars correspond to minimum and maximum average node distance for each cell size bin. At least 100 cells were analyzed for each condition. (C) Representative images of wild type and *pdk1*Δ cells expressing Mid1-mNG. mNG images are maximum intensity projections through thetop half of the cell with inverted fluorescence. Scale bar, 7 μm. (D) Quantification of average Mid1-mNeonGreen node distance versus cell size in wild type and *pdk1Δ* cells. Error bars correspond to minimum and maximum average node distance for each cell size bin. At least 100 cells were analyzed for each condition. All statistical tests performed in this figure are Welch’s unpaired t-tests. ns indicates *p* value > 0.05; * indicates *p* value < 0.05.

Finally, because cytokinetic rings failed to constrict in approximately 10% of *pdk1Δ* cells, we tested if Pdk1 contributes to cytokinesis by promoting signaling by the septation initiation network (SIN). The SIN pathway initiates cytokinetic ring constriction through the regulated acttivation and localization of Sid2 kinase to the SPB and cytokinetic ring (Simanis, 2015). We measured the timing of Sid2-GFP recruitment to the cytokinetic ring using SPB separation to normalize cytokinesis events between cells. We also tracked the cytokinetic ring marker Rlc1-mCherry in these same cells. Interestingly, we did not observe differences in the recruitment of Sid2 or Rlc1 when comparing wild type and *pdk1Δ* cells that completed cytokinesis (Figure 5A-C). Both the timing and levels of recruitment were unaffected. However, approximately 20% of *pdk1Δ* cells (4 out of 21) did not recruit Sid2 to the cytokinetic ring and subsequently failed cytokinesis (Figure 5A). In these *pdk1Δ* cells, Rlc1-mCherry formed a cytokinetic ring, but it did not initiate constriction and the cells did not complete cytokinesis. These *pdk1Δ* cells that failed cytokinesis also did not to recruit Sid2 to the SPB (Supplementary Figure S2F). This failure to recruit Sid2 to the SPB and cytokinetic ring was not seen in any wild type cells (0 out of 21). We conclude that SIN signaling fails in a subset of *pdk1Δ* cells, leading to binucleated cells and defects in later division events.

**Figure 5.**
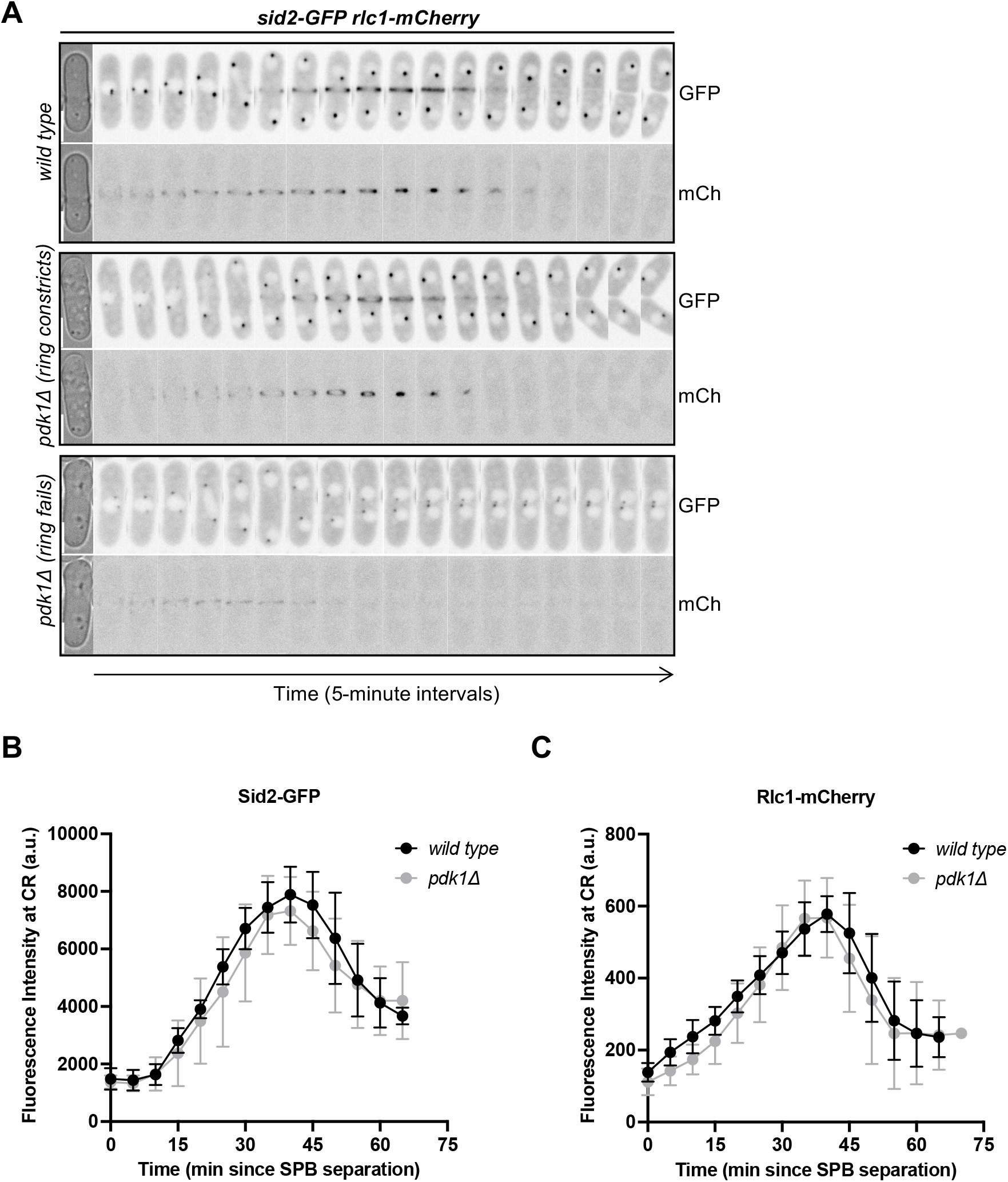
Localization of Sid2 kinase to the division site in *pdk1Δ* cells. (A) Time-lapse montage of wild type and *pdk1Δ* cells expressing Sid2-GFP and Rlc1-mCherry to mark the cytokinetic ring. Images are captured in 5-minute intervals. GFP and mCh images are sum projections of representative cells with inverted fluorescence. Brightfiled images are single middle focal planes from the first time point shown. Scale bar, 5 μm. (B) Quantification of Sid2-GFP recruitment to the cytokinetic ring in wild type and *pdk1Δ* cells. (C) Quantification of Rlc1-mCh recruitment to the cytokinetic ring in wild type and *pdk1Δ* cells. For panels B-C: time corresponds to minutes since Spindle Pole Body (SPB) separation as determined by Sid2-GFP signal; ti ≥ 10 cells each; and errors bars indicate mean and error ± standard deviation.

### Conclusions

We have identified new localization and functions for the conserved protein kinase Pdk1 in fission yeast. We found that Pdk1 localizes to and regulates cortical eisosomes, likely through phosphorylation of the core eisosome protein Pil1. Budding yeast Pdk1-related proteins play a similar role, but their function at eisosomes has been debated (Walther *et al*., 2007; Luo *et al*., 2008). Our results support a model where Pdk1 phosphorylates Pil1 to limit eisosome growth or stability. Future work will be needed to test this model *in vitro* with purified proteins. We also found that cytokinesis defects in *pdk1Δ* cells originate from abnormal spread of Mid1 nodes and failure to recruit Sid2. We did not observe localization of Pdk1 to cytokinetic nodes or the cytokinetic ring. Instead, it seems likely that Pdk1 activity at the SPB contributes to these cytokinesis-related events. In addition to linking Pdk1 with eisosomes and cytokinesis events, our results identify proteins that might be confirmed as Pdk1 substrates in future work. We note that many substrates of PDK-related kinases have a PDK1-interacting fragment (PIF), defined by the sequence motif F-X-X-F/Y-S/T/D-F/Y (Balendran *et al*., 1999; Biondi *et al*., 2000). We identified a PIF motif in Sid2 as well as the SIN protein Cut11. Both Sid2 and Cut11 localize at the SPB like Pdk1. We also found PIFs in the cytokinesis proteins Myp2 and Rga7. *In vitro* kinase assays with purified proteins will be needed to test if these PIF-containing proteins represent direct substrates of Pdk1. Future work on these proteins might define the molecular links between Pdk1 and cytokinesis. More generally, continued work on the role of Pdk1 in fission yeast cytokinesis has the potential to define the broad roles of PDK family kinases in eukaryotic cell growth and division.

## MATERIALS AND METHODS

### Strain construction and growth

Standard *Schizosaccharomyces pombe* media and methods were used (Moreno *et al*., 1991). Strains used in this study are listed in Supplementary Table S1. Gene deletions and tagging were performed using PCR and homologous recombination (Bähler *et al*., 1998). Pdk1-mNeonGreen was judged to be functional because the *pdk1-mNG* strain does not display any defect associated with *pdk1Δ* mutant. Pil1-mCherry is functional as judged by the lack of synthetic genetic interactions displayed by *pil1Δ* mutant (Kabeche *et al*., 2014).

*pDC99-PPdk1-Pdk1-mNeonGreen-Tadh1* was generated by Gibson assembly using PCR products. The Ppdk1-mNeonGreen insert was amplified from genomic DNA from JM8217 (*pdk1-mNeonGreen::hphMX6*). The pDC99 backbone was amplified from pJM1512 (*pDC99-Parf6-Arf6-mRuby-Tadh1*). The resulting plasmid was linearized by PCR and integrated into the *leu1* locus of JM737 (*ura4-D18*). Correct colonies were those that grew on EMM-Uri (uridine) medium but not on EMM-Leu (leucine) medium. The resulting strain was crossed to JM8289 (*pdk1Δ::kanMX6 ura4-D18 leu+*) to eliminate the copy at the endogenous *pdk1* locus.

In Supplementary Figure S1, double mutants were constructed by crossing JM8220 (*pdk1Δ*::*kanMX6*) to known cytokinetic mutants, including *rng2-D5* and *cdc15-140*. In Supplementary Figure S1C, cells were spotted by 10-fold serial dilutions on YE4S plates and incubated at 25° C, 30° C, 32° C, or 34° C for three days before imaging. Cells were grown in YE4S at 25° C before being stained with Blankophor to mark the cell wall in Supplementary Figure S1D.

### Microscopy and Image Analysis

All imaging was done at room temperature using a Nikon Eclipse Ti-2E microscope with a Yokogawa CSU-W1 spinning disk confocal system. This system was equipped with a Photometrics Prime BSI sCMOS camera; a 100x 1.45-NA CFI60 Plan Apochromat Lambda objective; and 405-, 488-, and 561 nm laser lines. mNG/GFP/YFP proteins were imaged using ET525/36m emission filter, and mCherry proteins were imaged using 605/52m emission filter. Cells were grown in YE4S at 25° C unless otherwise stated.

All image analysis was performed in ImageJ (Schneider *et al*., 2012). All statistics and graphs were generated using Prism10 (GraphPad software). Colocalization between Pdk1 and Rlc1 puncta were performed as previously described (Opalko *et al*., 2022). In short, the ImageJ plug-in comdetv5.5 was used to identify spots of Pdk1 and Rlc1. The distance between spots was set to 3 pixels, pixel size was set to 3, and intensity threshold was 3 for both channels. The low frequency of overlapping Pdk1 and Rlc1 spots (∽15% in Supplementary Figure S1B) contrasts strong colocalization (∽75%) seen for other proteins in the same Rlc1-containing structures from past studies (Opalko *et al*., 2022).

All strains imaged in Figure 1 were grown in in YE4S (yeast extract with 4 supplements) at 25° C. In Figure 1A (left), Z stacks were obtained by acquiring 13 slices with a 0.5 μm step size through the entire cell. mNG images are maximum intensity projections with inverted fluorescence. In Figure 1A (right), single middle focal planes and maximum intensity projections are shown of a single cell expressing Pdk1-mNeonGreen. Wild type cells expressing Pdk1-mNeonGreen, Rlc1-mCherry to mark the cytokinetic ring, and Sad1-mCherry to label the Spindle Pole Body (SPB) were imaged under glass coverslips in Figure 1B. Z stacks were obtained by acquiring 13 slices with a 0.5 μm step size through the entire cell. mCh and mNG images are maximum intensity projections through the top half of the cell. Merged images are maximum intensity projections of mCh (magenta) and mNG (cyan) channels overlaid. In Figure 1C, wild type cells expressing Pdk1-mNeonGreen and Pil1-mCherry to mark eisosomes were imaged under glass coverslips. Z stacks were obtained by acquiring 13 slices with a 0.5 μm step size through the entire cell. mCh and mNG images are maximum intensity projections with inverted fluorescence. Merged images are maximum intensity projections of mCh (magenta) and mNG (cyan) channels overlaid. Representative wild type and *pil1Δ* cells expressing Pdk1-mNeonGreen were imaged under glass coverslips in Figure 1D. Z stacks were obtained by acquiring 13 slices with a 0.5 μm step size through the entire cell. Left panel, mNG images are maximum intensity projections with inverted fluorescence. Right panel, single middle focal planes and maximum intensity projections for a single wild type and *pil1Δ* cell expressing Pdk1-mNeonGreen. All brightfield images represent a single middle focal plane.

Wild type, *pdk1Δ*, and *2x-pdk1* cells expressing Pil1-mCherry were grown in YE4S at 25° C in Figure 2A. The cells were maintained in rich media for 48 hours prior to being imaged under glass coverslips. In Figure 2A, Z stacks were obtained by acquiring 13 slices with a 0.5 μm step size through the entire cell. mCh images are maximum intensity projections with inverted fluorescence. Brightfield images are single middle focal planes. To measure eisosome length in Figure 2B, maximum intensity projections through the top half of the cell were generated. Individual Pil1-mCherry filaments were measured using the straight-line tool in Image J. Prior to being stained with Blankophor to mark the cell wall, the strains in Figure 2D were grown in YE4S at 25° C for 48 hours. Single middle focal plane images with inverted fluorescence are shown in Figure 2D.

In Figure 3A, wild type and *pdk1*Δ cells expressing Rlc1-mNeonGreen to mark the contractile actomyosin ring (CAR) and Sad1-meGFP to label the SPB were imaged on YE4S agarose pads. Representative sum projections with inverted fluorescence are shown. Brightfield images correspond to the first time point captured. Images were captured in 5-minute intervals. Z stacks were obtained by acquiring 7 slices with a 0.5 μm step size through half the cell. The onset of mitosis was monitored by Sad1-meGFP signal (Nabeshima *et al*., 1998). In Figure 3B, ring assembly was defined as the time taken to form an intact cytokinetic ring from the onset of mitosis. In Figure 3C, ring maturation was reported as the period between cytokinetic ring formation and the onset of ring constriction. In Figure 3D, ring constriction was described as the time from the initiation of ring constriction to the disappearance of Rlc1-mNeonGreen from the division site.

In Figure 4A and Figure 4C, wild type and *pdk1*Δ cells expressing (A) Cdr2-meGFP and (C) Mid1-mNeonGreen were grown in YE4S at 25° C for 48 hours. Cells were imaged in YE4S and mounted under glass coverslips. Z stacks were obtained by acquiring 13 slices with 0.5 μm step size through the entire cell. Brightfield images are a single middle focal plane. mEGFP and mNG images are maximum intensity projections with inverted fluorescence. To determine the average node distance of Cdr2-mEGFP (Figure 4B) and Mid1-mNeonGreen (Figure 4D), a maximum intensity projection through the top half of the cell was generated and the length of the medial node domain was measured for both sides of the cell. The resulting values were averaged and plotted against the length of the cell examined.

In Figure 5, wild type and *pdk1Δ* cells expressing Sid2-GFP and Rlc1-mCherry were imaged on YE4S agarose pads in 5-minute intervals. Z stacks were obtained by acquiring 7 slices with a 0.5 μm step size through half the cell. The onset of mitosis was monitored by Spindle Pole Body separation using Sid2-GFP signal (Nabeshima *et al*., 1998). In Figure 5A, brightfield images are a single middle focal plane corresponding to the first time point analyzed. GFP and mCh images are sum projections with inverted fluorescence. To determine the fluorescence intensity of Sid2-GFP and Rlc1-mCherry at the cytokinetic ring, an ROI was drawn around the ring using the rectangular tool in ImageJ. This ROI was used to measure the integrated density from sum intensity projections. The same ROI was moved to a neighboring region without any cells to measure the integrated density of the background. This value was used to subtract background signal. This process was repeated at all time points examined. The mean fluorescence intensity of Sid2-GFP (Figure 5B) and Rlc1-mCherry (Figure 5C) were plotted against time since Spindle Pole Body separation.

The time-lapse in Figure 5 was further analyzed to examine the recruitment of Sid2-GFP to the SPBs in both wild type and *pdk1Δ* cells. The latter was further categorized into two groups: those in which the cytokinetic ring successfully constricted and those in which the cytokinetic ring failed to constrict. An ROI was drawn around both SPBs using the freehand tool in ImageJ. This ROI was used to measure the integrated density from sum intensity projections. The same ROI was moved to a neighboring region without any cells to measure the integrated density of the background. This process was repeated at all time points examined. The sum fluorescence intensity of Sid2-GFP at the SPB was determined by adding the background-subtracted integrated density of both SPBs at each time point. The sum fluorescence intensity of Sid2-GFP at the SPB was plotted against time since SPB separation (Supplementary Figure S2F).

To determine the fluorescence intensity of different cytokinetic ring proteins in wild type and *pdk1*Δ cells, cells were grown in YE4S at 25° C for 48 hours. Cells were imaged in YE4S and mounted under glass coverslips. Z stacks were obtained by acquiring 13 slices with 0.5 μm step size through the entire cell. The rectangular tool in ImageJ was used to draw an ROI around the cytokinetic ring. This ROI was used to measure the integrated density from sum intensity projections. The same ROI was moved to a neighboring region without any cells to measure the integrated density. This value was used to subtract background signal.

### Western blots

Fission yeast whole cell lysates were prepared by growing cells in YE4S to mid-logarithmic phase. Two OD600 samples were harvested and rapidly frozen in liquid nitrogen. Cell pellets were resuspended in sample buffer (15% glycerol, 4.5% SDS, 97.5 mM Tris pH 6.8, 10% 2-mercaptoethanol, 50 mM β-glycerophosphate, 50 mM sodium fluoride, 5 mM sodium orthovanadate, 1x EDTA-free protease inhibitor cocktail (Sigma Aldrich). The samples were lysed with acid-washed glass beads (Sigma Aldrich) in a Mini-BeadBeater-16 (BioSpec, Bartlesville, OK) for 2 minutes at 4°C. Following this, cell lysates were incubated at 99° C for 5 minutes. Cell lysates were briefly centrifuged, and the supernatant was isolated as the clarified lysate. 8.0% acrylamide gels were poured containing 50 μM Phosbind acrylamide (APExBIO Catalog No. F4002) and 100μM MnCl2. 5µl of the cell lysates were separated by SDS-PAGE at 20 mAmps. Gels were washed once in transfer buffer containing 10 mM EDTA for 10 minutes. Next, gels were washed twice in transfer buffer without EDTA for 10 minutes. Blots were transferred to a nitrocellulose membrane using the Trans-blot Turbo Transfer System (Bio-Rad). Blots were first probed with custom anti-Pil1 primary antibodies at 4°C. Blots were washed in 1X TBST and then probed with goat anti-rabbit secondary antibody (LI-COR Lot No. C50331-05). Before being developed on a LiCor Odyssey CLx, blots were washed multiple times in 1X TBST and once in 1X TBS.

## ACKNOWLEDGEMENTS

We thank members of the Moseley lab for helpful discussions. This work was supported by a grant from the National Institutes of General Medical Sciences to J.B.M. (R35GM149248). This work was also supported by bioMT through NIH NIGMS grant P20GM113132.

---

**Supplementary Figure 1. Genetic interactions between *pdk1Δ* and cytokinesis mutants.** (A) Domain layout of the indicated PDK1-related protein kinases. (B) Quantification of Pdk1-mNG colocalization with Rlc1-mCherry. Datapoints represent percentage of Pdk1-mNG cortical puncta that colocalize with Rlc1-mCherry puncta in individual cells, as described in Materials and Methods. Bars represent mean ± SD; n = 19 cells. (C) Serial-dilution growth assays for each indicated strain at 25°, 30°, 32°, and 34° C. (D) Representative images of cells stained with Blankophor to mark the cell wall for the indicated strains. Images are a single middle focal plane with inverted fluorescence. Cells were grown at 25°. Scale bar, 14 μm. (E) Quantification of septation index for the indicated strains.

**Supplementary Figure 2.** (A-E) Quantification of the fluorescence intensity of the indicated proteins at the cytokinetic ring in wild type and *pdk1Δ* cells. ns indicates *p* value > 0.05; * indicates *p* value < 0.05; ** indicates *p* value < 0.005. N ≥ 25 cells each. All statistical tests performed in this figure are Welch’s unpaired t-tests. (F) Quantification of Sid2-GFP localization at the SPB for the indicated cell types.

## Notes

### Competing Interest Statement

The authors have declared no competing interest.

### Summary of Updates

This version of the manuscript has been revised with new data in Figures 2A-B, S1B, and S2F. Additional edits have been made in the text and figures.

