## Supplemental Table S1 for "Fission yeast Pdk1 kinase regulates cytokinesis and eisosomes"

| Strain | Description | Source | Figure |
|---|---|---|---|
| JM8215 | *pdk1-mNeonGreen::hphMX6 h-* | This study | 1A |
| JM8266 | *rlc1-mCherry::kanMX6 sad1-mCherry::natMX6 pdk1-mNeonGreen::hphMX6* | This study | 1B |
| JM8243 | *pdk1-mNeonGreen::hphMX6 pil1-mCherry::natMX6* | This study | 1C |
| JM8217 | *pdk1-mNeonGreen::hphMX6 h+* | This study | 1D |
| JM8254 | *pdk1-mNeonGreen::hphMX6 pil1Δ::natMX6 h-* | This study | 1D |
| JM1262 | *pil1-mCherry::natMX6 h-* | Lab Stock | 2A-B |
| JM8245 | *pil1-mCherry::natMX6 pdk1Δ::kanMX6* | This study | 2A-B |
| JM8674 | *pil1-mCherry::natMX6 ura4::pDC99-Ppdk1-pdk1-Tpdk1 h+* | This study | 2A-B |
| JM8269 | *ura4::pDC99-Ppdk1-pdk1-Tpdk1 ura4-D18 leu- h-* | This study | 2C |
| JM1243 | *pil1Δ::natMX6 h-* | Lab Stock | 2C |
| JM8219 | *pdk1Δ::kan h-* | This study | 2C-D |
| JM367 | *975 h+* | Lab Stock | 2C-D |
| JM8282 | *pdk1Δ::kanMX6 pil1Δ::natMX6* | This study | 2D |
| JM8257 | *rlc1-mNeonGreen::hphMX6 sad1-meGFP::kanMX6 pdk1Δ::kanMX6* | This study | 3A-D |
| JM5099 | *rlc1-mNeonGreen::hphMX6 sad1-meGFP::kanMX6 h-* | Lab Stock | 3A-D |
| JM6869 | *cdr2-mEGFP::kanMX6 h-* | Lab Stock | 4A-B |
| JM8296 | *cdr2-mEGFP::kanMX6 pdk1Δ::kanMX6* | This study | 4A-B |
| JM6994 | *mid1-mNeonGreen::hphMX6 h-* | Lab Stock | 4C-D |
| JM8267 | *mid1-mNeonGreen::hphMX6 pdk1Δ::kanMX6* | This study | 4C-D |
| JM8524 | *sid2-GFP::ura4+ rlc1-mCherry::kanMX6 ura4-D18 ade6-M21X* | This study | 5A-C |
| JM8545 | *sid2-GFP::ura4+ rlc1-mCherry::kanMX6 pdk1Δ::kanMX6 ura4-D18 ade6-M21X* | This study | 5A-C |
| JM8220 | *pdk1Δ::kan h+* | This study | S1B |
| JM366 | *972 h-* | Lab Stock | S1B |
| JM66 | *cdc4-31 h-* | Lab Stock | S1C-E |
| JM611 | *cdc15-140 h-* | Lab Stock | S1C-E |
| JM685 | *rng2-D5 ura4-D18 leu1-32 ade6-M21X h-* | Lab Stock | S1C-E |
| JM8474 | *pdk1Δ::kanMX6 cdc4-31* | This study | S1C-E |
| JM8475 | *pdk1Δ::kanMX6 cdc15-140* | This study | S1C-E |
| JM8476 | *pdk1Δ::kanMX6 cdc15-140* | This study | S1C-E |
| JM8477 | *pdk1Δ::kanMX6 rng2-D5* | This study | S1C-E |
| JM5024 | *rlc1-mNeonGreen::hphMX6 h-* | Lab Stock | S2A |
| JM8298 | *rlc1-mNeonGreen::hphMX6 pdk1Δ::kanMX6* | This study | S2A |
| JM99 | *myp2-mYFP::kanMX6 ade6-M210 leu1-32 ura4-D18 h-* | Lab Stock | S2B |
| JM8329 | *myp2-mYFP::kanMX6 pdk1Δ::kanMX6 ura4-D18* | This study | S2B |
| JM2191 | *rga7-meGFP::kanMX6 h-* | Lab Stock | S2C |
| JM8331 | *rga7-meGFP::kanMX6 pdk1Δ::kanMX6* | This study | S2C |
| JM8 | *kanMX6-Pmyo2-GFP-myo2 ade6-M210 leu1-32 ura4-D18 h+* | Lab Stock | S2D |
| JM8295 | *kanMX6-Pmyo2-GFP-myo2 pdk1Δ::kanMX6* | This study | S2D |
| JM7587 | *cdc12-mNeonGreen::hphMX6 h-* | Lab Stock | S2E |
| JM8297 | *cdc12-mNeonGreen::mNG::hphMX6 pdk1Δ::kanMX6* | This study | S2E |
