## Supplementary figures and images for "Fission yeast Pdk1 kinase regulates cytokinesis and eisosomes"

### Supplemental Figures S1-S2

#### Supplementary Figure 1.

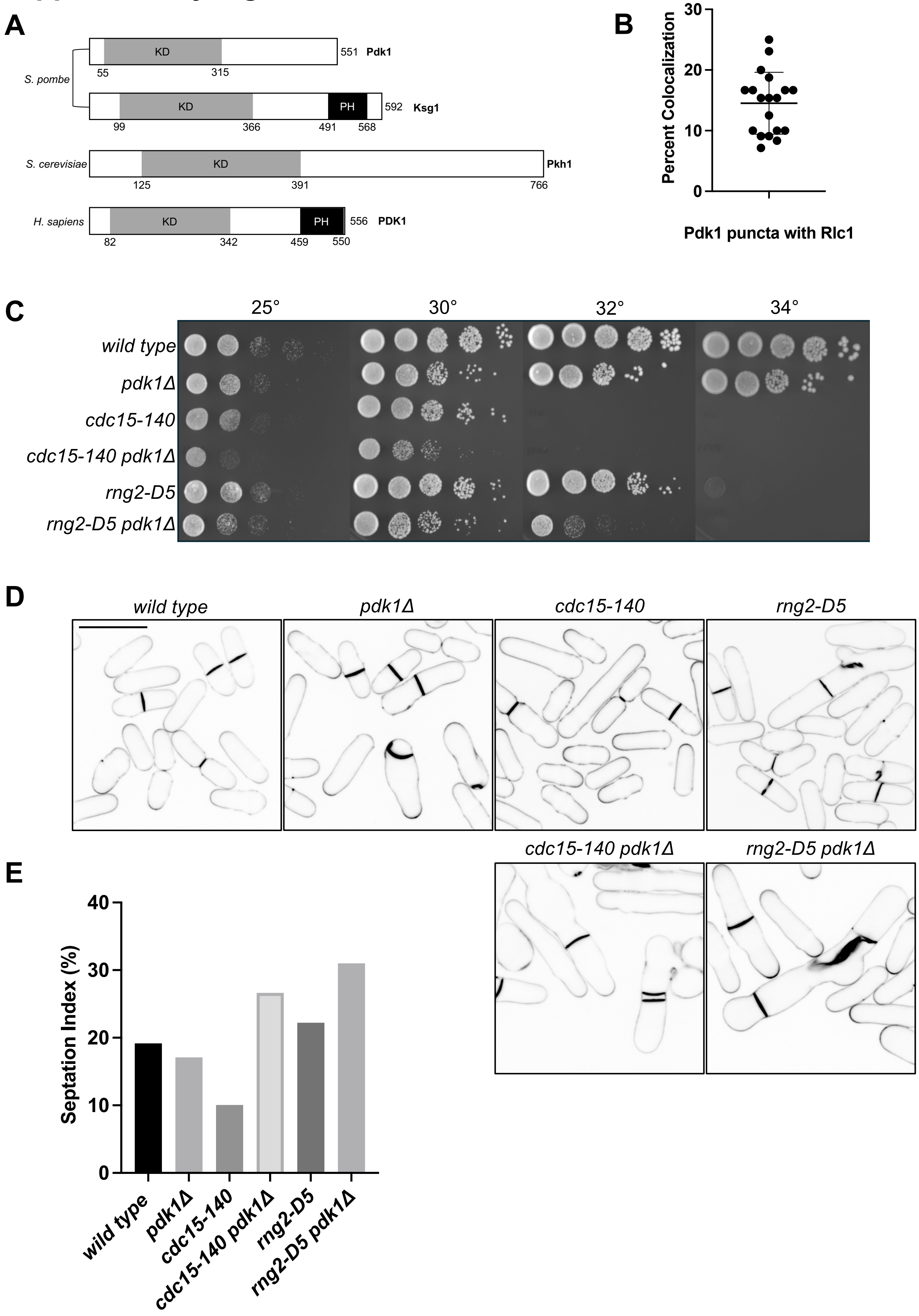

#### Supplementary Figure 2.

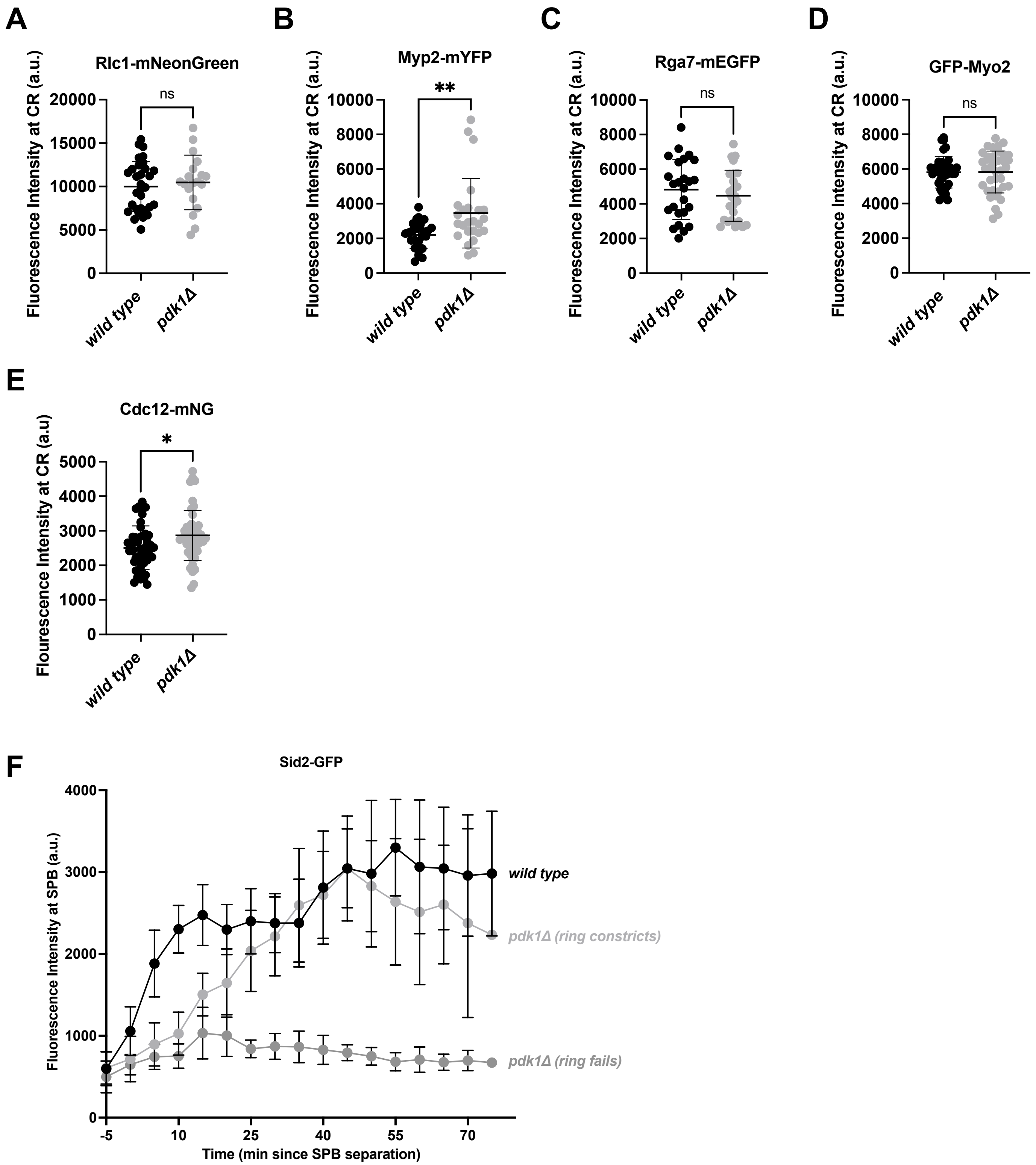
